# Associations between COVID-19 and putative markers of neuroinflammation: A diffusion basis spectrum imaging study

**DOI:** 10.1101/2023.07.20.549891

**Authors:** Wei Zhang, Aaron J Gorelik, Qing Wang, Sara A Norton, Tamara Hershey, Arpana Agrawal, Janine D Bijsterbosch, Ryan Bogdan

## Abstract

COVID-19 remains a significant international public health concern. Yet, the mechanisms through which symptomatology emerges remain poorly understood. While SARS-CoV-2 infection may induce prolonged inflammation within the central nervous system, the evidence primarily stems from limited small-scale case investigations. To address this gap, our study capitalized on longitudinal UK Biobank neuroimaging data acquired prior to and following COVID-19 testing (N=416 including n=224 COVID-19 cases; M_age_=58.6). Putative neuroinflammation was assessed in gray matter structures and white matter tracts using non-invasive Diffusion Basis Spectrum Imaging (DBSI), which estimates inflammation-related cellularity (DBSI-restricted fraction; DBSI-RF) and vasogenic edema (DBSI-hindered fraction; DBSI-HF).We hypothesized that COVID-19 case status would be associated with increases in DBSI markers after accounting for potential confound (age, sex, race, body mass index, smoking frequency, and data acquisition interval) and multiple testing.

COVID-19 case status was not significantly associated with DBSI-RF (|β|’s<0.28, p_FDR_ >0.05), but with greater DBSI-HF in left pre- and post-central gyri and right middle frontal gyrus (β’s>0.3, all p_FDR_=0.03). Intriguingly, the brain areas exhibiting increased putative vasogenic edema had previously been linked to COVID-19-related functional and structural alterations, whereas brain regions displaying subtle differences in cellularity between COVID-19 cases and controls included regions within or functionally connected to the olfactory network, which has been implicated in COVID-19 psychopathology.

Nevertheless, our study might not have captured acute and transitory neuroinflammatory effects linked to SARS-CoV-2 infection, possibly due to symptom resolution before the imaging scan. Future research is warranted to explore the potential time- and symptom-dependent neuroinflammatory relationship with COVID-19.

## Introduction

The ongoing COVID-19 pandemic remains a threat to global health and economies. In America alone, at least 42% of adults have been diagnosed with COVID-19 during their lifetime (1), and approximately 20% of them experience longer-term consequences, collectively referred to as long-COVID (2). These consequences include issues like loss of smell, psychopathology symptoms (e.g., sleep disorders, depression, anxiety), fatigue, cognitive impairment (also known as “brain fog”), and increased mortality (3–6). Concurrently, the pandemic continues to impose a substantial worldwide economic burden (7). Within the United Sates, the total estimated economic toll surged to $3.7 trillion USD in 2022 (8), up from the initial projections of $2.6 trillion USD (9). This encompasses expenses tied to diminished quality of life, decreased earnings, and escalated medical costs stemming from the long-lasting effects of COVID-19 (10, 11). These persistent challenges underscore the vital need for a comprehensive understanding of the underlying mechanisms through which COVID-19 impacts both health and behavior.

Accumulating evidence suggests that COVID-19 may have a profound impact on brain structure (6,12–14). Specifically, neurological events associated with COVID-19 such as stroke, vascular thrombosis, and microbleeds have been located throughout both cortical and deep subcortical structures (15), and individuals with SARS-Cov-2 infection have shown both cross-sectional and longitudinal changes in grey matter and white matter areas that are associated with cognitive impairment, sensory abnormalities, and mental health issues even months after the first infection (13, 14). For example, a recent study using longitudinal imaging data acquired prior to and following COVID-19 testing (total N=785 including n=401 COVID-19 cases), found reductions in cortical thickness and volume for individuals with a COVID-19 diagnosis in the orbitofrontal cortex and related regions (e.g., piriform cortex) that are involved in the olfactory network (16). Notably, these COVID-19 related longitudinal brain structure changes were also observed in individuals who had not been hospitalized (16). Evidence showing the association between COVID-19-related brain structural alterations and impaired cognitive performance further suggests that some COVID-19 related symptomatology may be attributable to changes in brain structure (21) and linked to elevated inflammation in the brain (22, 23). Interestingly, these brain-behavior associations may not resolve with time since COVID-19 related differences in brain structure have been observed one- and two-years following hospitalization (24, 25).

To elucidate the potential mechanisms contributing to these COVID-19-related structural changes in the brain, it becomes crucial to delve into the inflammatory processes in the central nervous system (26, 27). SARS-CoV-2 infection leads to activation of inflammatory signaling cascades, which can become dysregulated resulting in excessively high inflammation, known as cytokine storms (28–30). This hyperactivity of the immune system can disrupt the blood-brain barrier (31), enabling peripheral inflammatory markers to gain access into the CNS, which can directly increase neuroinflammation and may then impact brain structure (32). As such, neuroinflammation may occur in COVID-19 patients at the acute phase of disease (33–35) and persist in long-COVID patients with neuropsychiatric symptoms (22, 36). Brain olfactory regions may be particularly vulnerable to these hyperactive responses. For example, a study of golden hamsters found that SARS-CoV-2 infection may increase microglial and infiltrating macrophage activation in olfactory tissues, which was associated with behavioral alterations in scent-based food finding (37). Further evidence comes from post-mortem data in human COVID-19 patients and a rhesus macaque model of COVID-19 showing inflammation in the blood-brain barrier (i.e., choroid plexus; (38)), T-cell infiltration, and microglia activation (39).

While still in its preliminary stages, few positron emission tomography (PET) case studies have provided *in vivo* data establishing a connection between COVID-19 and neuroinflammation. These studies have demonstrated widespread increases in [^18^F]DPA-714 binding throughout the brain in two long-COVID patients (36), higher levels of translocator protein (TSPO) in the brainstem that were correlated with the clinical progression of one patient who had experienced both COVID-19 vaccination and subsequent infection (40). Additionally, elevated translocator protein distribution volume (TSPO V_T_) was observed in 20 participants who continued to suffer from persistent depressive and cognitive symptoms after initially experiencing mild to moderate SARS-CoV-2 infection (22). However, as of now, there have yet to be any large-scale investigations of neuroinflammation in the context of COVID-19.

Here, we tested whether COVID-19 is associated with changes in putative neuroinflammation in a cohort of N=416 individuals (n=244 infected cases and n=192 non-infected controls) from the “COVID-19 repeat imaging” sub-study of the UK Biobank (41). This sub-study collected neuroimaging data from individuals prior to and following a COVID-19 positive or negative test (e.g., based on diagnoses comprising PCR test, hospital impatient admission or GP records, as well as home-based test; see details in source of positive test result below in Methods). Neuroinflammation was assessed with putative markers of neuroinflammation-related cellularity and vasogenic edema. These markers were derived from diffusion-weighted imaging data, employing the Diffusion Basis Spectrum Imaging (DBSI) technique.

DBSI is an extension of standard Diffusion Tensor Imaging (DTI). While DTI focuses solely on estimating direction-dependent water movement parameters through anisotropic tensors, DBSI represents the diffusion-weighted imaging signal with multiple anisotropic and isotropic tensors (42–44). More importantly, the DBSI approach provides a spectrum of isotropic diffusion metrics, including restricted fraction (DBSI-RF), which indicates inflammation-related cellularity –either from cell proliferation or infiltration. Higher values of DBSI-RF suggest higher levels of inflammatory cell fraction. DBSI-RF as a putative neuroinflammation marker has been validated in a series of experiments. It demonstrated associations with inflammation-related cellularity derived from immunohistochemistry in an experimental mouse model of induced autoimmune encephalomyelitis (44), and with stain-quantitated nuclei and microglia density (i.e., cellularity) from post-mortem human brain tissues (45). Higher levels of DBSI-RF have also been linked to inflammation-related conditions, including multiple sclerosis (43,46–48), obesity (49, 50), HIV (51), Alzheimer’s disease (45,49,52), and depression (49), as well as markers of disease progression (45, 54). In addition to DBSI-RF, hindered fraction from the DBSI estimation (DBSI-HF) has shown promise as a putative marker for indicating inflammation-related cerebral edema (43, 55), and has been linked to inflammatory conditions including obesity (49, 50), and Alzheimer Disease (52). Nevertheless, both DBSI-RF and DBSI-HF as proposed markers of neuroinflammation would benefit from more validation work in these condition to which it is applied.

Here, employing this novel and non-invasive approach to assess neuroinflammation, we hypothesized that SARS-CoV-2 infection would be associated with increases in DBSI neuroinflammation markers across the brain with the most profound differences in brain regions that showed the strongest COVID-19 related structural changes (e.g., orbitofrontal cortex, piriform cortex) in the human brain (20)) or neuroinflammatory changes (e.g., microglia activation in olfactory bulb) in post-mortem or animal studies (37,39,56).

## Materials and Methods

### Sample

The UK Biobank (UKB) is a large-scale study (N > 500,000 participants) designed to examine the genetic, environmental, biological, and behavioral correlates of broad-spectrum health outcomes and related phenotypes (58). In February 2021, a UKB sub-study, the ‘COVID-19 repeat imaging study,’ was launched to collect neuroimaging data at a second timepoint, following either a positive (cases) or negative (controls) COVID-19 test, among individuals who completed a neuroimaging session prior to the COVID-19 pandemic to study longitudinal neuroimaging correlates of SARS-CoV-2 infection. COVID-19 positivity/negativity prior to the second neuroimaging session was determined from 3 sources: 1) hospital records contained in the Hospital Episode Statistics (a database containing admissions, outpatient appointments, and attendances and emergency admissions to English National Health Service Hospitals), or 2) primary care (GP) data, and 3) a record of a positive COVID-19 antibody test obtained from a home-based lateral flow kit sent to participants. For individuals who completed home testing and were vaccinated, a second testing kit was collected to ensure that any antibodies detected were from infection as opposed to recent vaccination (20). Participants were identified as COVID-19 positive cases if they had a positive test record on any of these data sources. COVID-19 negative participants (i.e., controls) were then selected in this sub-study from the remaining participants that were previously imaged before the pandemic to achieve 1:1 matching to the COVID-19 positive cases on sex, race (white/non-white), age (date of birth), location and date of the first imaging assessment. Details of inclusion criteria and case-control matching are provided in online documentation (https://biobank.ndph.ox.ac.uk/showcase/showcase/docs/casecontrol_covidimaging.pdf).

Data from the ‘COVID-19 repeat imaging study’ has been released on a rolling basis. As of March 24, 2023, we identified N = 416 (including n = 224 COVID-19 positive case) participants from the matched case-control list (variable ID 41000 in the UKB Data Showcase), who met the following inclusion criteria of the current study: 1) no mismatch between self-reported and genetic sexes, and 2) no missing data in any of the measures used in the current study. Demographic characteristics of the present study sample are summarized in **Table 1** and consistent with study design, were comparable between COVID-19 case and control participants. We also identified COVID-19 testing date for n=219 (97.77%) of the COVID-19 cases (UKB variable ID 40100). For this group of participants, the COVID-19 testing preceded the second imaging session by 128.3±70.8 days on average (range 37-372 days). Although unfortunately, data for COVID-19 related symptomatology and vaccination status were unavailable for the current study, we identified a small subgroup of COVID-19 case participants (n=13; 5.80%) who had a record for hospital inpatient admissions (variable ID 41001 in the UKB Data Showcase). Demographic information for hospitalized vs. non-hospitalized cases is summarized in **Supplemental Table S1**.

**Table 1.**
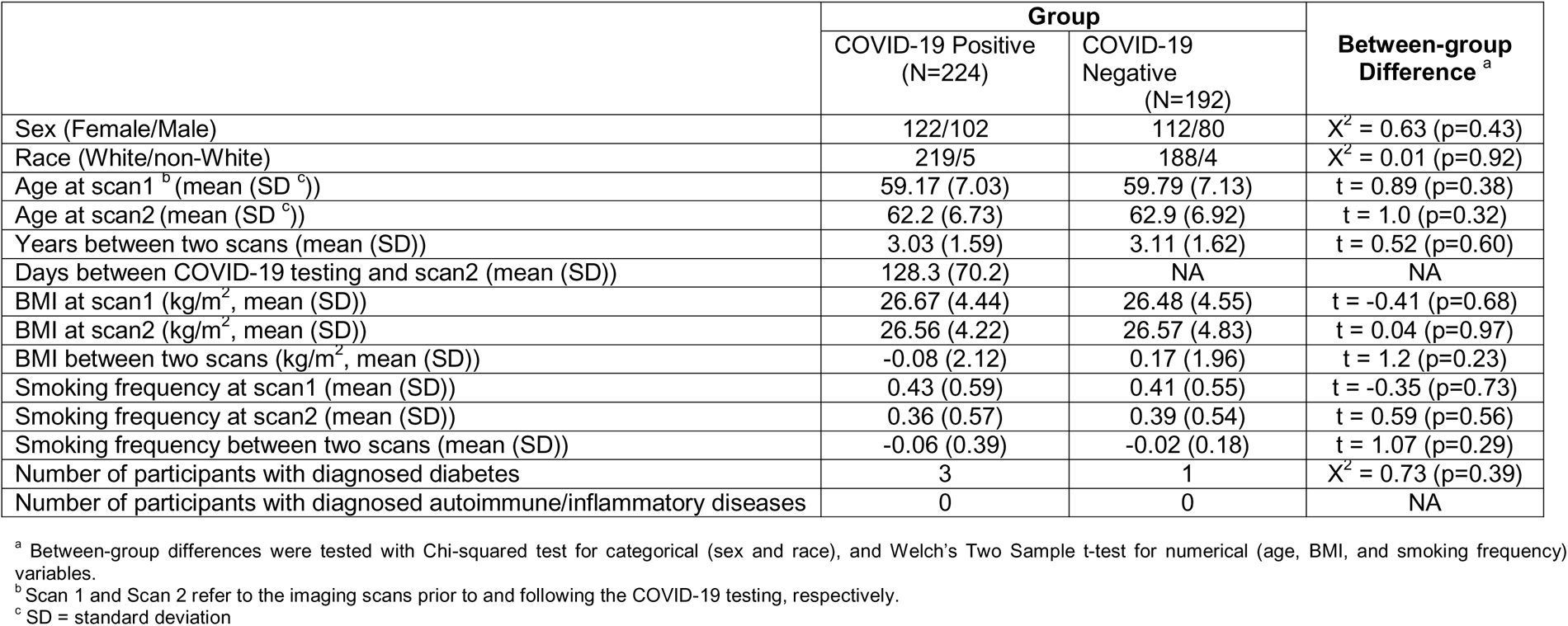
Demographics of the Study Sample.

|  | Group |  | Between-group Difference <sup>a</sup> |
| --- | --- | --- | --- |
|  | COVID-19 Positive (N=224) | COVID-19 Negative (N=192) |  |
| Sex (Female/Male) | 122/102 | 112/80 | $\chi^2 = 0.63$ (p=0.43) |
| Race (White/non-White) | 219/5 | 188/4 | $\chi^2 = 0.01$ (p=0.92) |
| Age at scan1 <sup>b</sup> (mean (SD <sup>c</sup>)) | 59.17 (7.03) | 59.79 (7.13) | t = 0.89 (p=0.38) |
| Age at scan2 (mean (SD <sup>c</sup>)) | 62.2 (6.73) | 62.9 (6.92) | t = 1.0 (p=0.32) |
| Years between two scans (mean (SD)) | 3.03 (1.59) | 3.11 (1.62) | t = 0.52 (p=0.60) |
| Days between COVID-19 testing and scan2 (mean (SD)) | 128.3 (70.2) | NA | NA |
| BMI at scan1 (kg/m<sup>2</sup>, mean (SD)) | 26.67 (4.44) | 26.48 (4.55) | t = -0.41 (p=0.68) |
| BMI at scan2 (kg/m<sup>2</sup>, mean (SD)) | 26.56 (4.22) | 26.57 (4.83) | t = 0.04 (p=0.97) |
| BMI between two scans (kg/m<sup>2</sup>, mean (SD)) | -0.08 (2.12) | 0.17 (1.96) | t = 1.2 (p=0.23) |
| Smoking frequency at scan1 (mean (SD)) | 0.43 (0.59) | 0.41 (0.55) | t = -0.35 (p=0.73) |
| Smoking frequency at scan2 (mean (SD)) | 0.36 (0.57) | 0.39 (0.54) | t = 0.59 (p=0.56) |
| Smoking frequency between two scans (mean (SD)) | -0.06 (0.39) | -0.02 (0.18) | t = 1.07 (p=0.29) |
| Number of participants with diagnosed diabetes | 3 | 1 | $\chi^2 = 0.73$ (p=0.39) |
| Number of participants with diagnosed autoimmune/inflammatory diseases | 0 | 0 | NA |

<sup>a</sup> Between-group differences were tested with Chi-squared test for categorical (sex and race), and Welch's Two Sample t-test for numerical (age, BMI, and smoking frequency) variables.
<sup>b</sup> Scan 1 and Scan 2 refer to the imaging scans prior to and following the COVID-19 testing, respectively.
<sup>c</sup> SD = standard deviation

### Imaging Acquisition and Processing

Diffusion Weighted Imaging (DWI) and T1-weighted structural MRI scans that were processed by the UKB Brain Imaging group (50) were used in the present study. Briefly, DWI data were acquired using a two-shell approach (b_1_ = 1000 and b_2_ = 2000 s/mm2) with 50 distinct diffusion-encoding directions within each shell. This multi-shell acquisition scheme is comparable to those used for the development of DBSI (42, 52). In general, multi-shell diffusion sequences have advantages of reduced sensitivity to the confounding effects of in-scanner motion (59) and have been shown to improve the estimation for free water corrected measures and free water fraction (60), and the angular resolution of orientation distribution functions (61). In addition, the UKB employed EPI-based spin echo acquisitions with opposite phase encode direction to reduce image distortion while reducing acquisition time (41), further increasing signal sensitivity. The acquired DWI data were then preprocessed to correct for eddy currents, head motion, outlier slice, and gradient distortion. The preprocessed data are available for download from the UKB database. T1 structural MRI data were acquired using an in-plane acceleration sequence and preprocessed to remove the skull and corrected for gradient distortion. Further processing on the T1 images was then carried out using FreeSurfer software, which produced images, surface files, and summary outputs, all available for download from the UKB database (https://biobank.ndph.ox.ac.uk/showcase/). More information about the acquisition protocols, image processing pipeline, and derived imaging measures can be found in the UKLBiobankLImagingLDocumentation (https://biobank.ctsu.ox.ac.uk/crystal/crystal/docs/brain_mri.pdf) and studies by Alfaro-Almagro et al. (62) and Miller et al. (41) .

### Diffusion Basis Spectrum Imaging and Neuroinflammation Indices

The DBSI technique employs a linear combination of anisotropic and isotropic tensors in describing the diffusion-weighted imaging signal, thereby improving sensitivity and specificity of estimated diffusion property (37, 42, 50). The primary DBSI neuroinflammation marker, known as the restricted (cellular) fraction (DBSI-RF), correlates with elevated cellularity and has been associated with activated microglia and astrogliosis in conditions such as multiple sclerosis (43,46,48,57,63) and Alzheimer’s disease (45,52,64). DBSI hindered (extracellular) fraction (DBSI-HF), which is indicative of vasogenic edema, has also been linked to neuroinflammation (65, 66) and inflammatory conditions including obesity (49, 50) and Alzheimer Disease (52).

To indicate neuroinflammation levels for specific brain structures, we applied the DBSI analysis package that was developed in house using the MATLAB (42) to the DWI data and used pre-defined brain structures (i.e., regions of interest, ROIs) as masks to extract region-specific neuroinflammation indices. Specifically, we used gray matter parcellations generated by the Automatic Subcortical Segmentation (67) and Desikan-Killiany cortical atlases (68). The resulting gray matter ROIs consisted of 14 bilateral subcortical and 66 cortical parcellations, representing n = 7 and n = 33 subcortical and cortical structures, respectively. Additionally, n = 20 white matter tracts from both left and right hemispheres were extracted from a probabilistic tractography atlas (JHU-ICBM-tracts) with the lowest probability of 25% at a given brain voxel (69). We used this probability threshold to ensure that each individual white matter tract could be identified in the subject-specific diffusion images (i.e., increasing probability would result in uneven tract numbers identified at the individual level).

To extract neuroinflammation indices per individual for each of the gray and white matter ROIs, we first created individual-specific ROI masks by registering T1 structural (i.e., FreeSurfer outputs) and MNI standard diffusion images (i.e., white matter tracts) with native diffusion images. The mean DBSI-RF and DBSI-HF values across all voxels within each ROI were then calculated for each individual participant, respectively. As more than half of the entire gray matter structures (n = 21) showed a correlation less than 0.6 between the left and right hemispheric homologues for the baseline DBSI-RF values (i.e., pre-COVID), we considered each parcellation as an independent ROI for statistical analysis (i.e., not combined across hemispheres). Similarly, all 20 white matter tracts were considered as separate ROI as the left-right correlations for all tracts from the baseline DBSI-RF values were below 0.6. We applied the same pipeline to DBSI-HF and included each individual gray matter parcellation and white matter tract as a unique ROI in the subsequent analyses involving DBSI-HF, due to low baseline correlations (r<0.06) between two hemispheric homologues for many gray matter structures (n=20) and white matter tracts (n=10).

#### Covariates

Following a prior study (20), we included genetic sex (UKB variable ID 22001), ethnicity (UKB variable ID 21000), as well as differences between pre- and post-COVID assessments in age (UKB variable ID 21003) as covariates in this study. We further included changes in body mass index (BMI; UKB variable ID 21001), in smoking status (UKB variable ID 20116), and in date (i.e., number of days; UKB variable ID 53) between two assessments to adjust for potential confounds that may contribute to the changes in neuroinflammation from pre-to post-COVID. As in this prior study (20), we also used white versus non-white for ethnicity in all models.

#### Statistical Analysis

We tested whether COVID-19 cases differed from controls on neuroinflammation, as indexed by DBSI-RF. Mirroring the analytic strategy of a prior study linking COVID-19 case status to brain structural changes (20), we conducted a series of linear regressions in which COVID-19 case/control status was modeled as a predictor of post-COVID regional neuroinflammation, while accounting for pre-COVID neuroinflammation and the covariates described above. Separate models were conducted for each individual ROI and false discovery rate (FDR) was applied to adjust for multiple testing within gray and white matter ROIs, separately (i.e., 40 tests for gray matter ROIs, and 20 tests for white matter ROIs). This analytic pipeline was repeated for our primary (DBSI-RF) and secondary (DBSI-HF) neuroinflammation markers, respectively.

As it is plausible that neuroinflammation may reflect a transient phenomenon resolving over time, we further conducted post-hoc analyses on data of participants whose positive COVID-19 test occurred ≤ 60 (n = 75) and ≤90 (n = 23) days prior to the neuroimaging session and conducted group comparisons of each of these two subgroups with controls separately. Furthermore, we repeated the main analyses without the restriction on the assessment time interval after excluding the data from the COVID-19 positive cases with a hospitalization record (N=13) to explore the potential impact of symptom severity on the association between neuroinflammation and COVID-19.

As changes in putative neuroinflammation within each ROI may not occur uniformly, we further explored the association between COVID-19 and whole-brain voxel-wise neuroinflammation, as indexed by DBSI-RF. To this end, we first registered all participants’ DBSI-RF maps (i.e., both pre- and post-COVID maps) with the standard MNI brain template and obtained a differential map between two scans per individual (i.e., scan 2 minus scan 1; delta DBSI-map in MNI space). Using these delta maps as input, we then conducted permutation tests (permuted n = 5000) with FSL *randomise (70)* and a Threshold-Free Cluster Enhancement (TFCE) method (71) to identify delta DBSI-RF “clusters” in the brain that differ between COVID-19 negative and positive individuals (i.e., case-minus-control contrast and control-minus-case contrast), while accounting for covariates (i.e., sex, ethnicity, data acquisition interval, and changes in age, BMI, and smoking status). Categorical variables (e.g., sex, group) were dummy coded before permutation testing. These post-hoc and exploratory analyses were conducted for DBSI-RF and DBSI-HF, separately.

## Results

After multiple testing correction, COVID-19 was not associated with any changes in DBSI-RF values across all gray matter regions and white matter tracts. There were only a handful of nominally significant associations (i.e., uncorrected p<0.05; **Figure 1**; **Table 2**). Gray matter regions included the left caudal middle frontal gyrus (β = -0.14, p_uncorrected_ = 0.048), superior parietal lobule (β = -0.14, p_uncorrected_ = 0.047), and postcentral gyrus (β = -0.14, p_uncorrected_ = 0.041), as well as the right caudate (β = 0.18, p_uncorrected_ = 0.039), amygdala (β = 0.22, p_uncorrected_ = 0.013), caudal anterior cingulate cortex (β=0.25, p_uncorrected_ = 0.003), rostral anterior cingulate cortex (β = 0.27, p_uncorrected_ = 0.003), lateral orbitofrontal cortex (β=0.19, p_uncorrected_ = 0.037), and fusiform gyrus (β = 0.19, p_uncorrected_ = 0.029). Only one region was nominally significant in white matter: the temporal proportion of the left superior longitudinal fasciculus (β = 0.25, p_uncorrected_ = 0.008). When we repeated these analyses excluding COVID-19 case participants who had been hospitalized at the time of COVID-19 testing, results remained similar with negligibly small changes in beta values (see **Supplemental Table S2**).

**Figure 1.**
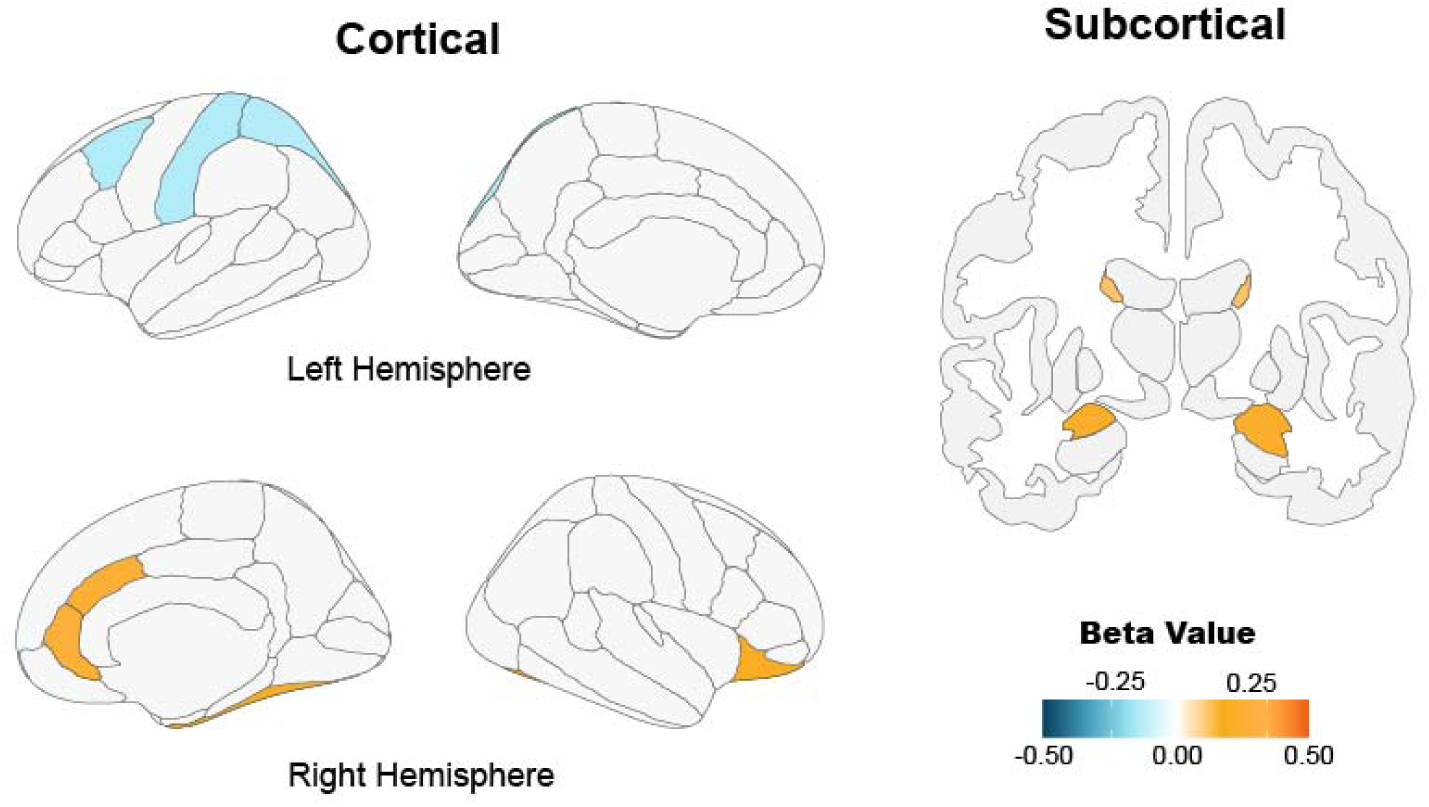
Gray Matter Regions with Nominal Increase in DBSI-RF (p_uncorrected_<0.05).

In contrast, COVID-19 was significantly associated with DBSI-HF values in the pre- and post-central gyri in the left hemisphere, as well as the caudal proportion of the right middle frontal gyrus after FDR corrections (β’s > 0.3, FDR p’s = 0.03; **Figure 2**). In addition, a set of cortical regions and white matter tracts exhibited nominally significant associations, such as bilateral superior frontal gyrus, bilateral pars triangularis, and right cingulum bundle (β’s > 0.15, uncorrected p’s < 0.05; see full list in **Table 3**). These results remained when excluding the hospitalized participants (**Supplemental Table S3**).

**Figure 2.**
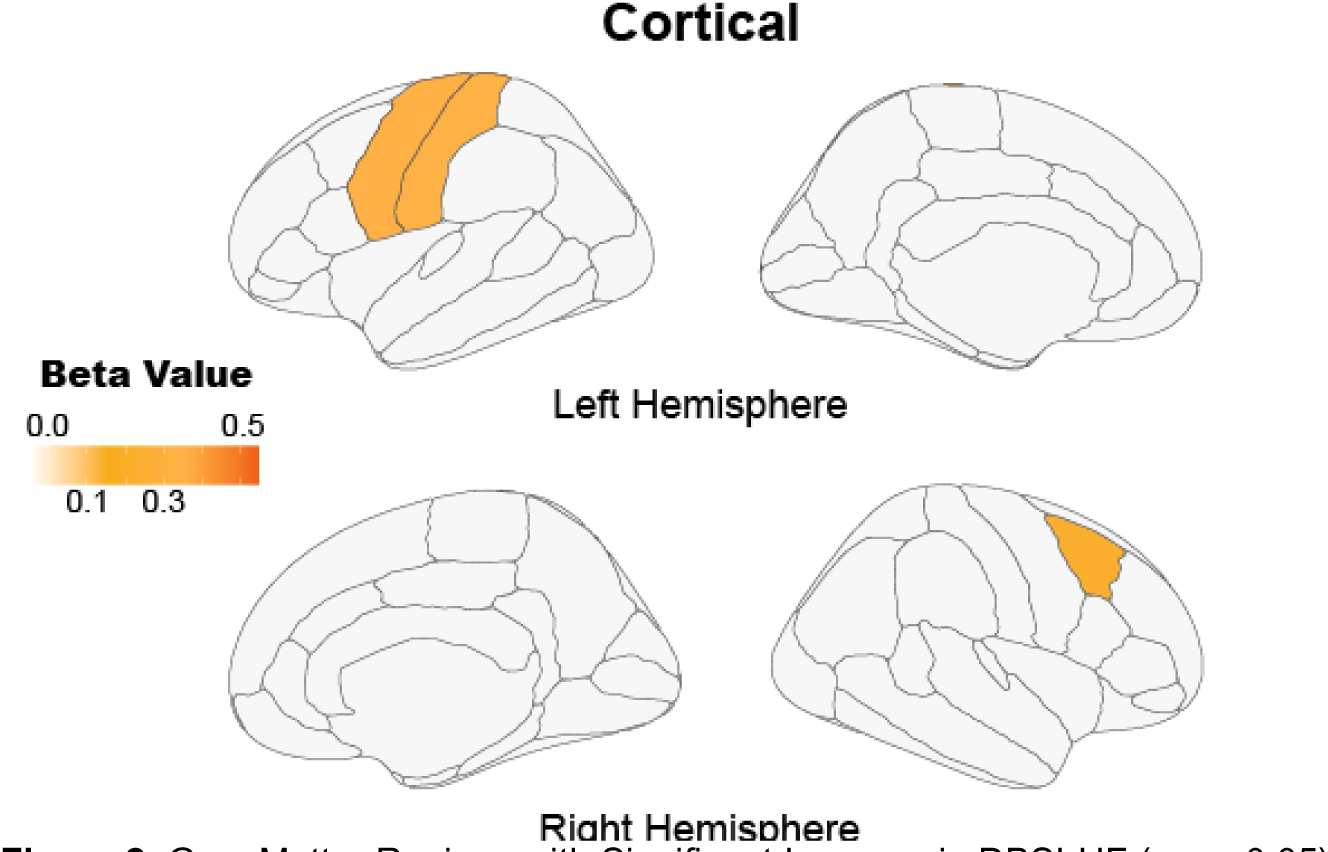
Gray Matter Regions with Significant Increase in DBSI-HF (p_FDR_<0.05).

**Table 2.** Nominally significant between-group differences in DBSI-RF.

| Brain Structure | ROI (L/R)<sup>a</sup> | Statistics<sup>b</sup> |  |  |  |
| --- | --- | --- | --- | --- | --- |
|  |  | B<br>[95% CI]<sup>c</sup> | *t* | Uncorrected<br>P-value | FDR<br>corrected<br>P-value<sup>d</sup> |
| Subcortical |  |  |  |  |  |
|  | Caudate (R) | 0.18<br>[0.01, 0.35] | 2.08 | 0.039 | 0.424 |
|  | Amygdala (R) | 0.22<br>[0.04, 0.41] | 2.49 | 0.013 | 0.356 |
| Cortical |  |  |  |  |  |
|  | Caudal Middle Frontal (L) | -0.14<br>[-0.28, 0.001] | -1.99 | 0.048 | 0.424 |
|  | Postcentral (L) | -0.14<br>[-0.27, 0.003] | -2.05 | 0.041 | 0.424 |
|  | Superior Parietal (L) | -0.13<br>[-.027, 0.001] | -1.99 | 0.047 | 0.424 |
|  | Caudal Anterior Cingulate (R) | 0.25<br>[0.05, 0.44] | 2.54 | 0.011 | 0.356 |
|  | Fusiform (R) | 0.19<br>[0.02, 0.36] | 2.20 | 0.029 | 0.424 |
|  | Lateral Orbitofrontal (R) | 0.19<br>[0.01, 0.38] | 2.09 | 0.037 | 0.424 |
|  | Rostral Anterior Cingulate (R) | 0.27<br>[0.09, 0.46] | 2.94 | 0.003 | 0.279 |
| White Matter Tracts |  |  |  |  |  |
|  | Superior Longitudinal Fasciculus (L; Temporal Part) | 0.25<br>[0.06, 0.44] | 2.66 | 0.008 | 0.165 |

<sup>a</sup> ROI = Regions of Interest; L/R = Left / Right hemisphere<sup>b</sup> Group difference between the COVID-19 positive and negative participants after adjusting for the covariates.<sup>c</sup> CI = Confidence Interval<sup>d</sup> FDR = False Discovery Rate

**Table 3.** Nominally and statistically significant between-group differences in DBSI-HF.

| Brain Structure | ROI (L/R) <sup>a</sup> | Statistics <sup>b</sup> |  |  |  |
| --- | --- | --- | --- | --- | --- |
|  |  | B<br>[95% CI] <sup>c</sup> | *t* | Uncorrected<br>P-value | FDR<br>corrected<br>P-value <sup>d</sup> |
| Cortical |  |  |  |  |  |
|  | Caudal Middle Frontal (L) | 0.28<br>[0.08,0.48] | 2.75 | 0.006 | 0.074 |
|  | Inferior Parietal (L) | 0.20<br>[0.004,0.39] | 2.0 | 0.046 | 0.174 |
|  | Middle Temporal (L) | 0.22<br>[0.04,0.40] | 2.34 | 0.02 | 0.145 |
|  | Paracentral (L) | 0.26<br>[0.07,0.45 ] | 2.74 | 0.006 | 0.074 |
|  | Pars Opercularis (L) | 0.21<br>[0.02,0.40] | 2.18 | 0.03 | 0.16 |
|  | Pars Triangularis (L) | 0.25<br>[0.06,0.45] | 2.54 | 0.01 | 0.102 |
|  | Postcentral (L) | 0.32<br>[0.13,0.51] | 3.28 | 0.001 | **0.03\*** |
|  | Precentral (L) | 0.33<br>[0.14,0.53] | 3.38 | 0.0008 | **0.03\*** |
|  | Rostral Middle Frontal (L) | 0.27<br>[0.07,0.46] | 2.68 | 0.008 | 0.077 |
|  | Superior Frontal (L) | 0.28<br>[0.08,0.47] | 2.82 | 0.005 | 0.074 |
|  | Superior Parietal (L) | 0.21<br>[0.02,0.40] | 2.12 | 0.03 | 0.160 |
|  | Supramarginal (L) | 0.26<br>[0.08,0.45] | 2.78 | 0.006 | 0.074 |
|  | Transverse Temporal | 0.18<br>[0.01,0.35] | 2.08 | 0.038 | 0.16 |
|  | Insula (L) | 0.20<br>[0.006,0.39] | 2.02 | 0.044 | 0.174 |
|  | Caudal Middle Frontal (R) | 0.33<br>[0.13,0.52] | 3.30 | 0.001 | **0.03\*** |
|  | Isthmus Cingulate (R) | 0.20<br>[0.02,0.39] | 2.16 | 0.03 | 0.16 |
|  | Pars Triangularis (R) | 0.23<br>[0.03,0.43] | 2.28 | 0.02 | 0.155 |
|  | Precentral (R) | 0.21<br>[0.01,0.40] | 2.09 | 0.037 | 0.16 |
|  | Precuneus (R) | 0.20<br>[0.01,0.38] | 2.10 | 0.036 | 0.16 |
|  | Rostral Middle Frontal (R) | 0.24<br>[0.05,0.44] | 2.41 | 0.016 | 0.13 |
|  | Superior Frontal (R) | 0.22<br>[0.02,0.42] | 2.16 | 0.03 | 0.16 |
| White Matter Tracts |  |  |  |  |  |
|  | Cingulum (R; Anterior and Superior Portion) | 0.19<br>[0.04,0.34] | 2.54 | 0.011 | 0.138 |
|  | Superior Longitudinal Fasciculus (R; Temporal Part) | -0.21<br>[-0.38,-0.04] | -2.47 | 0.014 | 0.138 |

<sup>a</sup> ROI = Regions of Interest; L/R = Left / Right hemisphere
<sup>b</sup> Group difference between the COVID-19 positive and negative participants after adjusting for the covariates.
<sup>c</sup> CI = Confidence Interval
<sup>d</sup> FDR = False Discovery Rate
\* Significant results after FDR corrections

### Proximity of Scan to COVID-19 Diagnosis

After accounting for multiple testing, we did not observe significant differences in either DBSI-RF or DBSI-HF values between participants whose COVID-19 positive tests were closer in time to the neuroimaging session (i.e., ≤ 60 or ≤ 90 days following COVID-19 test) and non-infected control participants. Yet, nominally higher DBSI-RF values were observed in several brain structures and tracts, including rostral anterior cingulate cortex in the right hemisphere (β’s > 0.3, uncorrected p’s < 0.05), which were also observed in the full sample (**Table 4**). Further, DBSI-HF values in several regions including the left pars triangularis, right cuneus cortex, and the rostral portion of the anterior cingulate cortex showed nominally significant associations with COVID-19 in both subsets across different time-windows (|β|’s > 0.2, uncorrected p’s < 0.05; **Table 5**).

### Whole-brain voxel-wise differences in putative neuroinflammation

After applying threshold-free cluster enhancement (TFCE), we did not observe any clusters that reached a statistical significance (all p’s>0.05) for DBSI-RF or DBSI-HF.

**Table 4.** Nominally significant between-group differences in DBSI-RF in potentially acute COVID-19 cases.

| Brain Structure | ROI (L/R) <sup>a</sup> | Statistics |  |  |  |  |  |  |  |
| --- | --- | --- | --- | --- | --- | --- | --- | --- | --- |
|  |  | ≤ 60 Days <sup>b</sup> |  |  |  | ≤ 90 Days <sup>b</sup> |  |  |  |
| | | $\beta$<br>[95% CI] <sup>c</sup> | *t* | Uncorrected<br>P-value | FDR<br>corrected<br>P-value | $\beta$<br>[95% CI] <sup>c</sup> | *t* | Uncorrected<br>P-value | FDR<br>corrected P-value <sup>d</sup> |
| Subcortical |  |  |  |  |  |  |  |  |  |
|  | Hippocampus (R) |  |  |  |  | 0.27<br>[0.03, 0.52] | 2.22 | 0.027 | 0.463 |
| Cortical |  |  |  |  |  |  |  |  |  |
|  | Pars Triangularis (L) | -0.39<br>[-0.75, -0.02] | -2.11 | 0.036 | 0.597 | -0.24<br>[-0.46, -0.02] | -2.22 | 0.027 | 0.463 |
|  | Postcentral (L) |  |  |  |  | -0.21<br>[-0.41, -0.02] | -2.19 | 0.029 | 0.463 |
|  | Cuneus (R) | 0.37<br>[0.02, 0.72] | 2.13 | 0.035 | 0.597 | 0.23<br>[0.01, 0.45] | 2.12 | 0.035 | 0.463 |
|  | Fusiform (R) |  |  |  |  | 0.28<br>[0.04, 0.53] | 2.33 | 0.020 | 0.463 |
|  | Pericalcarine Cortex (R) | 0.41<br>[0.02, 0.80] | 2.10 | 0.037 | 0.597 |  |  |  |  |
|  | Rostral Anterior Cingulate (R) | 0.52<br>[0.09, 0.95] | 2.40 | 0.011 | 0.597 | 0.32<br>[0.06, 0.58] | 2.45 | 0.015 | 0.463 |
|  | Rostral Middle Frontal (R) | -0.41<br>[-0.76, 0.07] | -2.39 | 0.029 | 0.597 |  |  |  |  |
| White Matter Tracts |  |  |  |  |  |  |  |  |  |
|  | Superior Longitudinal Fasciculus (L; Temporal Part) |  |  |  |  | 0.34<br>[0.07, 0.60] | 2.53 | 0.012 | 0.243 |

<sup>a</sup> ROI = Regions of Interest; L/R = Left / Right hemisphere
<sup>b</sup> Group comparisons of COVID-19 negative individuals with COVID-19 positive individuals whose COVID-19 positive testing occurred within 60 or 90 days of their second imaging scan.
<sup>c</sup> CI = Confidence Interval
<sup>d</sup> FDR = False Discovery Rate

**Table 5.** Nominally significant between-group differences in DBSI-HF in potentially acute COVID-19 cases.

| Brain Structure | ROI (L/R) <sup>a</sup> | Statistics |  |  |  |  |  |  |  |
| --- | --- | --- | --- | --- | --- | --- | --- | --- | --- |
|  |  | ≤ 60 Days <sup>b</sup> |  |  |  | ≤ 90 Days <sup>b</sup> |  |  |  |
| | | $\beta$<br>[95% CI] <sup>c</sup> | *t* | Uncorrected<br>P-value | FDR<br>corrected<br>P-value | $\beta$<br>[95% CI] <sup>c</sup> | *t* | Uncorrected<br>P-value | FDR<br>corrected P-value <sup>d</sup> |
| Subcortical |  |  |  |  |  |  |  |  |  |
|  | Thalamus (L) |  |  |  |  | 0.30<br>[0.09,0.50] | 2.81 | 0.005 | 0.065 |
| Cortical |  |  |  |  |  |  |  |  |  |
|  | Caudal Anterior Cingulate (L) |  |  |  |  | 0.28<br>[0.02,0.54] | 2.07 | 0.04 | 0.177 |
|  | Cuneus (L) |  |  |  |  | 0.28<br>[0.02, 0.53] | 2.15 | 0.033 | 0.162 |
|  | Inferior Parietal (L) |  |  |  |  | 0.34<br>[0.07,0.60] | 2.46 | 0.01 | 0.13 |
|  | Middle Temporal (L) |  |  |  |  | 0.41<br>[0.15,0.67] | 3.12 | 0.002 | 0.065 |
|  | Postcentral (L) |  |  |  |  | 0.38<br>[0.12,0.65] | 2.84 | 0.005 | 0.065 |
|  | Precentral (L) |  |  |  |  | 0.39<br>[0.12,0.66] | 2.84 | 0.005 | 0.065 |
|  | Rostral Middle Frontal (L) |  |  |  |  | 0.33<br>[0.06,0.60] | 2.37 | 0.02 | 0.143 |
|  | Superior Frontal (L) |  |  |  |  | 0.28<br>[0.01,0.55] | 2.07 | 0.04 | 0.177 |
|  | Superior Parietal (L) |  |  |  |  | 0.31<br>[0.04,0.58] | 2.24 | 0.026 | 0.149 |
|  | Supramarginal (L) |  |  |  |  | 0.39<br>[0.14,0.64] | 3.01 | 0.003 | 0.065 |
|  | Transverse Temporal (L) |  |  |  |  | 0.25<br>[0.02,0.48] | 2.17 | 0.03 | 0.162 |
|  | Caudal Middle Frontal (R) |  |  |  |  | 0.39<br>[0.13,0.66] | 2.87 | 0.004 | 0.065 |
|  | Cuneus (R) | 0.43<br>[0.05, 0.86] | 1.98 | 0.05 | 0.823 | 0.37<br>[0.11,0.63] | 2.79 | 0.006 | 0.065 |
|  | Isthmus Cingulate (R) |  |  |  |  | 0.30<br>[0.04,0.55] | 2.30 | 0.02 | 0.143 |
|  | Precentral (R) |  |  |  |  | 0.31<br>[0.05,0.58] | 2.33 | 0.02 | 0.143 |
|  | Precuneus (R) |  |  |  |  | 0.33<br>[0.08,0.59] | 2.55 | 0.01 | 0.114 |
|  | Superior Parietal (R) |  |  |  |  | 0.30<br>[0.04,0.56] | 2.28 | 0.02 | 0.143 |
| **White Matter Tracts** |  |  |  |  |  |  |  |  |  |
|  | Superior Longitudinal Fasciculus (L; Temporal Part) |  |  |  |  | 0.34<br>[0.07, 0.60] | 2.53 | 0.012 | 0.243 |
|  | Cingulum (R; Anterior and Superior Portion) | -0.44<br>[-0.86,-0.02] | -2.05 | 0.04 | 0.412 | -0.27 | -2.06 | 0.04 | 0.27 |
|  | Inferior Frontooccipital Fasciculus (R) | -0.41<br>[-0.79,-0.03] | -2.12 | 0.04 | 0.412 |  |  |  |  |
|  | Inferior Longitudinal Fasciculus (R) |  |  |  |  | -0.26<br>[-0.48,-0.04] | -2.28 | 0.02 | 0.237 |
|  | Superior Longitudinal Fasciculus (R; Temporal Part) |  |  |  |  | -0.32<br>[-0.57,-0.08] | -2.63 | 0.009 | 0.183 |

<sup>a</sup> ROI = Regions of Interest; L/R = Left / Right hemisphere
<sup>b</sup> Group comparisons of COVID-19 negative individuals with COVID-19 positive individuals whose COVID-19 positive testing occurred within 60 or 90 days of their second imaging scan.
<sup>c</sup> CI = Confidence Interval
<sup>d</sup> FDR = False Discovery Rate

## Discussion

In this study we examined whether COVID-19 case status is associated with changes in neuroinflammation, as indexed by DBSI-RF (cellularity) and DBSI-HF (vasogenic edema), using a unique UKB prospective case-control cohort (n=224 cases; n=129 controls) that was scanned prior to and following COVID-19 testing. Contrary to our hypotheses and prior studies on post-mortem brain tissue (38, 56) and *in vivo* imaging of long-COVID cases (36), we found no significant association between COVID-19 status and changes in DBSI-RF after multiple testing correction. Notably, several brain regions such as the orbitofrontal cortex, anterior cingulate cortex, ventral striatum, which have demonstrated the most significant structural changes following COVID-19 (20), showed increases in DBSI-RF at nominal levels of significance among COVID-19 cases (**Table 2**). These findings were replicated in analyses focused on case participants who were scanned closer to their COVID-19 test date (≤ 60 or 90 days; **Table 4**) and when excluding n=13 hospitalized individuals (**Table S2**). However, caution is warranted in interpreting these nominal findings due to their failure to survive multiple testing correction, and the inconsistent directionality of findings across regions (e.g., reduced DBSI-RF among COVID-19 cases in the caudal middle frontal gyrus). Conversely, using DBSI-RF as a secondary measure for neuroinflammation, we observed significant associations between COVID-19 status and increased DBSI-HF values (**Table 3**). Although these findings persisted when excluding hospitalized individuals (see **Table S3**), they were no longer evident when examining COVID-19 case participants with a shorter time interval between COVID-19 testing and post-COVID imaging assessment (**Table 5**).

Together, our data suggest that the putative vasogenic edema may be more sensitive in capturing changes in neuroinflammatory processes following SARS-CoV-2 infection in our sample. Yet, it also remains possible that certain COVID-19 related neuroinflammatory processes such as neuroinflammation related to glia cell damage may be temporally constrained (e.g., functional recovery within 60-90 days after SARS-CoV-2 infection; (72–74)) and thus were resolved prior to the post-COVID scan.

### Infection-induced Neuroinflammation Progression and Clinical Heterogeneity

In contrast to the null findings of our primary neuroinflammation marker, DBSI-RF, other studies suggest that the acute infection of SARS-CoV-2 induces neuroinflammation, potentially leading to long-term consequences including neurological and psychiatric syndromes (75). Current evidence indicates that inflammation in the central nervous system may occur as early as several weeks after the onset of SARS-CoV-2 infection (33–35), and last up to two years after the infection with widespread influences in the brain in individuals suffering from long-COVID (22, 36). Alongside acute production of proinflammatory microglia (i.e., the most dominant immune cells in the brain) and a chronic loss of microglia (76), damages in astrocytes and oligodendrocytes were also observed in relation to COVID-19, which may undergo functional recovery 60-90 days after the infection onset, subsequently resulting in reductions of neuroinflammation (72–74). Alternatively, these glia cell damages may ultimately lead to neuronal cell death in the cerebral cortex (72, 77). These findings suggest that COVID-19-related inflammatory processes may diverge depending on the assessment timing (78). DBSI-RF, being an indirect proxy of inflammation-related cellularity from either immune cell proliferation or infiltration (42,44,46,52), raises the possibility that these inflammatory processes may have subsided by the time of the second imaging scan. Although DBSI-RF holds potential to capture a wide range of inflammatory processes, it may also require future investigation to determine its sensitivity in detecting inflammatory changes induced by SARS-CoV-2 infection in the central nervous system. Interestingly, while our primary neuroinflammation marker, DBSI-RF showed no association with COVID-19, we observed significant increases in the putative marker of vasogenic edema (i.e., DBSI-HF) for COVID-19 cases compared to non-infected controls. This result might indicate the presence of continued neuroinflammation in these case participants, consistent with the prolonged nature of neurological consequences following the resolution of acute COVID-19 illness. However, our mixed findings of DBSI markers may be influenced by the clinical heterogeneity of COVID-19 manifestations with some case participants possibly having fully recovered from the infection, while some others still experiencing some degrees of symptoms by the time of the second imaging assessment. The lack of information regarding the disease recovery and the real-time symptoms for COVID-19 positive cases adds complexity to our interpretation.

It should also be noted that the present findings were obtained after correcting for potential confounding effects of age, sex, ethnicity, BMI, smoking frequency, and data acquisition intervals. Additional factors may contribute to variations in clinical presentation and related neuroinflammation. For instance, a “two-hit” hypothesis on the link between microglial activation and COVID-19 severity suggests that predisposed conditions such as exposures to childhood trauma (i.e., first hit) may sensitize individuals’ microglia responses when facing the second immune challenge, such as COVID-19 (79). In line with this, a recent survey study has reported increased risks of developing post-infection conditions for individuals who had indicated prior-infection distress (80). Interestingly, however, psychological stress has also been linked to neuroinflammation in non-infected control participants during the COVID-19 pandemic, suggesting that neuroimmune activation may contribute to the development of symptoms not directly linked to the coronavirus SARS-CoV-2 (81). Future research may consider including stress-related factors to better understand neuroinflammation induced by SARS-CoV-2 infection and its implications.

### Nominal Associations of DBSI-RF with COVID-19

A few nominally significant results (i.e., before FDR correction) in DBSI-RF values are worth mentioning in the context of prior research findings. First, the lateral orbitofrontal cortex exhibited a nominally higher level of post-COVID neuroinflammation in individuals who were tested COVID-19 positive before the second imaging assessment. This finding aligns with prior animal studies that highlighted the susceptibility of the olfactory system, including the orbitofrontal regions, to SARS-CoV-2 virus invasion, showing the olfactory bulb as the primary entry point for the virus into the brain (82), with the nasal cavity’s olfactory epithelium identified as the enhanced binding site of the virus (83, 84). The initial virus invasion is then followed by a rapid and trans-neuronal spread of infection throughout the brain, including structures connected with the olfactory bulb as well as structures only remotely connected with the olfactory system (82). This wide spread infection may further manifest at the structural level of the brain such that even mildly or moderately infected individuals (i.e., non-hospitalized) exhibited greater reduction in grey matter thickness and tissue contrast in the orbitofrontal cortex (20) piriform cortex (i.e., the primary component of the olfactory network) including the amygdala, caudate, and the anterior cingulate cortex also showed nominally higher neuroinflammation levels in COVID-19 positive individuals (**Table 2**). Interestingly, these regions were previously reported with longitudinal anatomical changes in the same UK Biobank cohort who were COVID-19 positive with mild-to-moderate symptoms (20). In addition to these gray matter the temporal part of the superior longitudinal fasciculus (SLF), a white matter tract mainly connects the frontal and parietal cortices and plays a crucial role in language, attention, memory, and emotion (85, 86). Altered diffusion properties in SLF have been reported in post-COVID individuals who had mild to moderate acute COVID-19 (87, 88), and these alterations appear to be persistent long after recovery of COVID-19 (25, 87). Thus, the nominally elevated neuroinflammation in SLF might also be indicative of this pathological process induced by COVID-19. However, as these findings did not survive FDR correction, they should be interpreted with caution.

### Significant and Nominal Associations of DBSI-HF with COVID-19

While our primary measure of putative neuroinflammation-related cellularity did not exhibit significant associations with COVID-19, we observed an increased ratio of extracellular water (indicative of vasogenic edema) in COVID-19 cases compared to non-infected controls (**Table 3**). This is in line with previous observations of cerebral or brain tissue edema in patients with COVID-19 (21,89,90), Notably, heightened DBSI-HF in our study was found in primary motor and primary sensorimotor areas, which have previously shown aberrant connectivity patterns in individuals with acute SARS-CoV-2 infection (91), or six months after hospital discharge (92). Long-term changes in resting-state amplitude of low-frequency fluctuation (ALFF) were also reported for these areas in patients one year after recovery (16). Furthermore, the caudal middle frontal gyrus, exhibiting increased DBSI-HF in our study, has been linked to SARS-CoV-2-induced structural changes such as reduced cortical thickness (93). Interestingly, this COVID-19-related structural alteration within this brain region was further associated with increased inflammation markers in the cerebrospinal fluid (94). Yet, it is noteworthy that these prior findings were observed in individuals with a diverse spectrum of post-COVID sequelae (e.g., persistent fatigue and myalgia), suggesting that post-SARS-CoV-2 inflammatory processes may be symptom-dependent (95). Due to the lack of information about COVID-19 symptomatology in our study, future investigations are needed to explore the associations between COVID-19 symptoms and vasogenic edema present after SARS-CoV-2 infection.

### Limitations

It is important to consider study limitations when interpreting these data. First, COVID-19 positive cases in our study were defined solely by SARS-Cov-2 testing while symptom and severity assessments, as well as vaccination status were unavailable (96). This precluded us from investigating associations between the DBSI-derived putative neuroinflammation markers and COVID-19 symptomatology. While elevated peripheral inflammation markers have been observed in individuals with SARS-Cov-2 infection and are tied to the severity of COVID-19 symptoms (89,97–99), inflammatory responses appear to regress gradually during recovery in most patients (100, 101), This indicates a transient effect of inflammation that potentially is symptom dependent. It is therefore possible that our mixed findings in DBSI-RF and DBSI-HF are attributable to the study assessment schedule (i.e., second scan acquired on average 128 days post COVID-19) and lack of COVID-19 symptomatology data available to us. Future research is warranted to examine the impact of COVID-19 symptom severity on different neuroinflammatory processes following SARS-CoV-2 infection. Second, while the UK Biobank is population-based cohort, it is also self-selective (102) and predominantly White (**Table 1**), which may limit the generalizability of these findings. Third, information of COVID-19 testing date for control participants were unavailable, making it impossible to model within-subject variability in neuroinflammation changes due to differences in the time interval between COVID-19 testing and the second imaging assessment for the full study sample.

### Conclusion

While no statistical association was found between COVID-19 status and DBSI-RF, a relatively large sample of case participants (n = 224) demonstrated significant increases in DBSI-HF compared to non-infected controls (n = 192). These findings are consistent with prior research showing elevated neuroinflammation in postmortem brain tissues from severe COVID-19 patients (38, 77), and in long-COVID patients with persistent neurological or psychiatric symptoms (22, 36). Considering the potential impact of analysis timing on capturing distinct inflammatory responses to COVID-19 (78), our data suggest that neuroinflammation may be both time- and symptom-dependent. However, due to the lack of relevant information in our study, these findings should be interpreted with caution.

## Statement of Ethics

All participants in UK Biobank study have provided their written informed consent and that the current study has been granted an exemption from requiring ethics approval by the IRB at Washington University in St. Louis. This study was conducted under the UK Biobank Application ID 47267.

## Author Contributions

AA and RB conceived the idea. QW and TH contributed to DBSI processing and interpretation. WZ analyzed the data and wrote the first draft of the article together with AG. JB and RB contributed to supervision. All authors contributed to the manuscript preparation.

## Data Availability Statement

The UK Biobank data used in this study can be accessed by researchers upon application (https://www.ukbiobank.ac.uk/register-apply).

